# Elevated alpha diversity in disturbed sites obscures regional decline and homogenization of amphibian diversity

**DOI:** 10.1101/2021.09.21.461266

**Authors:** D. Matthias Dehling, J. Maximilian Dehling

**Author notes:** equal contribution.

## Abstract

Loss of natural habitat is one of the major threats for biodiversity worldwide. Habitat conversion not only changes diversity and species composition locally (alpha diversity) but might also lead to large-scale homogenization of species communities and decrease in regional species richness (gamma diversity). We investigated the effect of farmland conversion on amphibian communities in Rwanda and compared local and regional (country-wide) taxonomic, functional and phylogenetic diversity between natural and farmland sites (agricultural *marais*). Alpha diversity was higher in the disturbed farmland than in natural sites. However, species turnover among farmland sites was much lower than among natural sites, resulting in highly homogenized amphibian communities and much lower country-wide taxonomic, functional and phylogenetic gamma diversity in farmland compared to natural sites. The few frog species found in farmland were mostly disturbance-tolerant species that are widespread in Eastern Africa and beyond. In contrast, most of the regionally endemic frog species that make this region a continent-scale hotspot of amphibian diversity were found only in the natural habitats. Ongoing farmland conversion might lead to a loss of regional endemism and a widespread homogenization of species communities across sub-Saharan Africa.

## Introduction

Loss of native habitat due to anthropogenic land-cover change, such as deforestation and conversion into farmland, is one of the major drivers of species loss (Butchart et al. 2010; McGill et al. 2015), with severe impacts on global biodiversity (Bradshaw et al. 2009; Newbold et al. 2015). The loss of biodiversity has severe negative impacts on ecosystems functioning which, in turn, poses a threat to human well-being (Loreau et al. 2001; Hooper et al. 2005, 2012; Balvanera et al. 2006; Cardinale et al. 2012, Pasari 2013, Tilman et al. 2014, Murphy & Romanuk 2014; Johnson et al. 2017).

Disturbances such as habitat alterations might not only affect the number of species in a community (alpha diversity) but also their composition due to species replacements (de Coster 2015; Riemann et al. 2017). Hence, limiting the assessment of diversity on changes in alpha diversity— without considering changes in species composition—might underestimate the effect of habitat alterations on species communities (McGill et al. 2015). Moreover, species replacements do not only alter the composition of local species communities but can also lead to a homogenization of species communities at the regional scale, that is, across habitat types (van der Plas et al. 2016). No single species assemblage can simultaneously support all functions at high levels (Mori et al. 2018) and instead, sets of functions fulfilled by the species community from different habitat types complement each other on the regional scale (Hector & Bagchi 2007, Isbell et al. 2011), resulting in higher overall diversity of functions or “multifunctionality” (van der Plas et al. 2016). The functioning of ecosystems at high levels at the regional scale therefore requires diverse communities, with both high alpha and high beta diversity (Pasari et al. 2013, van der Plas et al. 2016, Dehling & Dehling 2021). The replacement of a species restricted to a certain habitat (habitat specialist) with widespread generalists leads to a loss of beta diversity between species communities and, consequently, to biotic homogenization at the regional scale (McKinney & Lockwood 1999). Although human-driven loss of beta diversity through biotic homogenization of communities could be as widespread and detrimental as the loss of alpha diversity (Murphy & Romanuk 2014; McGill et al. 2015; Newbold et al. 2015), studies on the effect of biotic homogenization on ecosystem functioning are still widely lacking (Loreau et al. 2003, Mori et al. 2018, Seibold et al. 2019, Felipe-Lucia et al. 2020).

The diversity of ecological assemblages in different habitat types has traditionally been measured as species richness, but increasingly, assessments of biological diversity are now considering the functional diversity and phylogenetic diversity of species assemblages. Functional diversity measures the combination in species traits that are associated with adaptations to the environment and species’ roles in ecological processes (Tilman 2001), and communities with higher functional alpha diversity provide a wider range of ecological functions (Cadotte et al. 2011; Flynn et al. 2011). Phylogenetic diversity measures the diversity of evolutionary lineages (Lean & Maclaurin 2016, Owen et al. 2019, Gumbs et al. 2020), which represent the adaptability of species to environmental changes (Brooks et al. 1992, Phillimore et al. 2007, Miraldo et al. 2016, Smith et al. 2017, Tucker et al. 2019). Since these different facets of diversity can idiosyncratically respond to disturbances (Flynn et al. 2009; Villéger et al. 2010; de Coster et al. 2015), consideration of taxonomic, functional and phylogenetic diversity might provide complementary insights into the effect of land-use change on species communities.

Amphibians are paramount for wetlands and aquatic habitats because they provide key ecological functions, especially in tropical regions (Gibbons et al. 2006, Hocking & Babbit 2014). Amphibian diversity worldwide is threatened by climatic change, invasive species and diseases (Beebee 1995; Kiesecker et al. 2001; Cheng et al. 2011), and especially by habitat loss (Wake & Vredenburg, 2008; Butchart et al. 2010). Despite the functional importance of amphibians, the effect of habitat alterations on amphibian communities is poorly understood: at the local scale, species richness in disturbed sites was found to be lower (Ernst & Rödel 2008; Gardner et al. 2007a,b; Gillespie et al. 2012; Angarita-M. et al. 2015; Jiménez-Robles et al. 2017), unchanged (Ernst et al. 2006, Oda et al. 2016), or even higher (Sinsch et al. 2012; Tumushimire et al. 2020) compared to natural sites. In addition, all studies so far have only focussed on changes in local alpha diversity, without considering the composition of amphibian communities. Most importantly, no study has investigated the effects of habitat alteration across scales, i.e. the extent to which the loss of natural habitats due to farmland conversion causes homogenization of amphibian communities and loss of regional gamma diversity and multifunctionality.

We investigated the effect of farmland conversion on the diversity and composition of amphibian communities in Rwanda. Rwanda is the most densely-populated country in continental Africa, with currently over 74 percent of the land surface being exploited for agriculture, and the remaining natural habitats threatened by growing demand for subsistence agriculture, livestock grazing, and fuel extraction (REMA 2017). Using data from 37 sites across the country, we compared the taxonomic (species richness), functional, and phylogenetic diversity of amphibian communities between natural sites (natural forest and savannah) and farmland (agricultural *marais*). We compared changes in the local diversity of amphibian communities (alpha diversity) with changes on the regional scale across Rwanda (gamma diversity). If amphibian communities in Rwanda responded positively to habitat alterations, as suggested by the high local species richness in agricultural marais (Sinsch et al. 2012; Tumushimire et al. 2020), we would expect similar levels of alpha, beta and gamma diversity in natural and farmland sites. However, if habitat alteration led to wide-scale species replacements, we would expect a homogenization of species communities in farmland sites at the regional scale, manifested in lower beta and gamma diversity compared to natural sites.

## Material and methods

### Sampling of amphibian communities

We sampled amphibian communities in natural habitats and farmland (agricultural marais) across Rwanda (Appendix1, Table S1). Natural habitats included swamps in natural forest with partial open areas surrounded by dense forest in Volcano National Park (NP), Gishwati-Mukura NP, and Nyungwe NP (“forest”, n = 11, 1804–3031 m a.s.l.) and wetlands and ephemeral ponds in natural savannah in Akagera NP (“savannah”, n = 11, 1287–1642 m a.s.l.). Farmland sites were distributed all over Rwanda and typically consisted of a mosaic of crop patches (< 0.1 ha) separated by small irrigation channels (n = 15, 1292–2348 m a.s.l.). We sampled each site six times at different times of the year, both at the beginning of the short rainy season (September, October) and at the maximum of the long rainy season (March, April). To assess seasonality of species activity, most sites—including all sites in Akagera NP, which are most strongly affected by seasonality—were also sampled during the short dry season (December, January) and at the beginning (May) and end (September) of the long dry season. We assessed the presence of species at each site through visual and acoustic encounter surveys (Rödel & Ernst 2004).

### Functional traits of frogs

For all frog species encountered, we collected ecological, morphological and life-history traits related to resource use and functional roles of species, following Ernst et al. (2006) and Cadotte et al. (2011): microhabitat use, calling site, breeding seasonality, egg deposition site and type, tadpole type, snout-vent length, head width, hind-limb length, hand and foot webbing, terminal disks (details in Appendix 1, Table S2). We obtained traits in the field as well as from specimens collected at the sites and from museum vouchers in the Royal Museum for Central Africa in Tervuren, Belgium. Additional life-history information was taken from Channing & Howell (2006).

### Amphibian phylogeny

For all frog species encountered, we compiled a phylogeny using the consensus tree from Jetz & Pyron (2018). Missing species (n = 3) were added at the tip of the most closely-related species.

### Comparison of alpha and gamma diversity

For each site, we determined the taxonomic, functional, and phylogenetic alpha diversity of amphibians. We measured taxonomic alpha diversity as species richness, the total number of frog species found at a site. We measured functional alpha diversity of amphibians as the diversity of their functional-trait combinations. We first calculated the pairwise differences in trait combinations between all frog species using Gower’s distance (since our functional traits included continuous and categorical data; Villéger et al. 2008). We then used non-metric multidimensional scaling to project all frog species into one common three-dimensional frog trait space where they were arranged according to the differences in their trait combinations. We calculated the functional-trait diversity for each site as functional richness, i.e. the volume of a convex hull in the three-dimensional trait space that includes all frog species found at that site (Villéger et al. 2008, Maire et al. 2015). We calculated phylogenetic diversity as Faith’s PD (Faith 1992), as the combined length of all branches that connect the frog species found at a site.

For each habitat category, we pooled the species across all sites to calculate the total species richness (taxonomic gamma diversity), total functional richness (functional gamma diversity), and total phylogenetic diversity (phylogenetic gamma diversity) found across Rwanda. For the farmland sites, we used all 15 sites sampled in farmland. For the natural sites, we sampled 1000 combinations of 7 and 8 sites from forest and savannah. To test whether our samples of frog communities were representative of the different habitat types (farmland, natural habitat), we calculated species-accumulation curves (Appendix 1, Fig. S1). Since some of the forest plots were located at higher elevations than the highest farmland sites, we repeated the comparison of alpha and gamma diversity without forest plots above 2500 m elevation (Appendix 2).

### Species and phylogenetic complementarity across sub-Saharan Africa

We used data from the IUCN to map the distribution of all amphibian species in sub-Saharan Africa (< 20° N) in 1° grid cells. We replaced the three grid cells for Rwanda with one grid cell that included the species found in natural habitats and two grid cells that included the species found in farmland (which is a conservative approach, as the farmland occupies a much larger area relative to natural habitat in Rwanda). For each grid cell, we calculated species richness and phylogenetic diversity of amphibians. In addition, we calculated the complementarity in species and phylogenetic diversity: the contribution of each grid cell to total species richness and phylogenetic diversity, measured relative to all other grid cells and relative to the species richness and diversity found in the grid cell (see Dehling et al. 2021 for functional complementarity). Complementarity is high for grid cells that include a high percentage of species with a small distribution, and it is low for grid cells that mostly include widespread species.

We used R 4.1 (R Core Team 2021) for all analyses.

## Results

### Comparison of alpha and gamma diversity

We recorded a total of 42 amphibian species at the study sites: 39 in natural habitats (22 in forest, 17 in savannah) and 19 species in farmland (Appendix 1, Table S1). Fifteen species (eleven savannah and four forest species) were shared between natural sites and farmland; three species (*Hyperolius lateralis*, *Ptychadena porosissima*, *P. uzungwensis*) were exclusively found in farmland.

Alpha diversity in natural habitats was lower than in farmland (*species richness*, natural sites: 6.5 ± 1.8, range 4–10, farmland: median ± SD 11 ± 3.2, range 6–17, Fig. 1a; *functional diversity*, natural sites: 1.26 ± 2.19, farmland: 2.24 ± 1.01, Fig. 1b; *phylogenetic diversity*, natural sites: 783 ± 191, farmland: 1248 ± 292, Fig. 1c). In contrast, gamma diversity was higher in natural habitats than in farmland (*species richness*, natural sites 35 ± 4, farmland 19, Fig. 1a; *functional diversity*, natural sites 20.8 ± 0.86, farmland 3.9, Fig. 1b; *phylogenetic diversity* natural sites 2501 ± 56, farmland 1731, Fig. 1c). The functional-trait combinations of amphibians in farmland were completely nested within those from natural sites (Fig. 2a). Most of the major phylogenetic lineages that were present in natural sites were still represented in the farmland, albeit by less than half of the branches (Fig. 2b,c). Species accumulation curves showed that our sampling of amphibian communities in farmland sites, but not in natural sites, reached saturation after 15 samples (Appendix 1, Fig. S1). The observed differences between natural sites and farmland are therefore conservative estimates and can be expected to be even more severe.

**Fig. 1.**
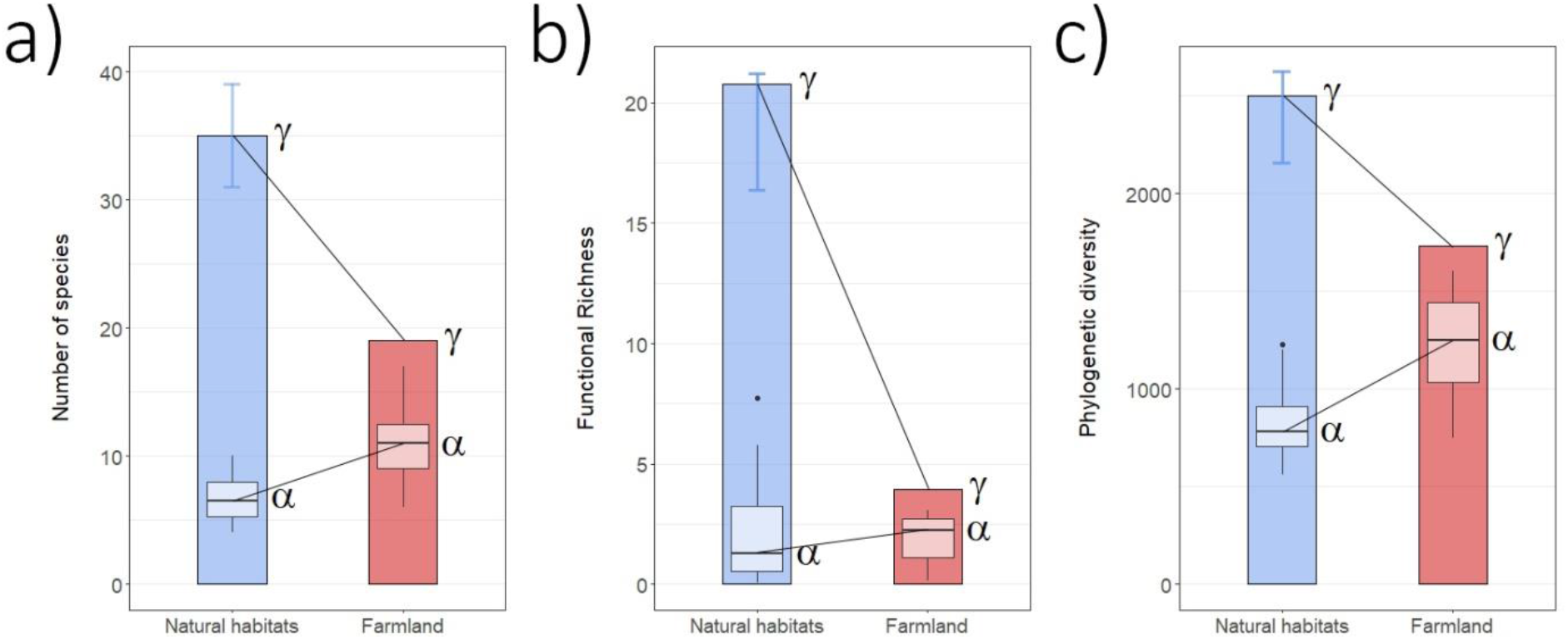
Alpha and gamma diversity of amphibian communities in natural habitats (natural forest, natural savannah; blue) vs farmland (red) across Rwanda. Diversity is measured as (a) taxonomic diversity (number of species), (b) functional diversity (functional richness), and (c) phylogenetic diversity (Faith’s PD) both on the level of local communities (alpha diversity) and once pooled across Rwanda (gamma diversity). Gamma diversity is shown as coloured bars; median and standard deviation of local alpha diversity (n = 15) are shown as lighter boxplots within bars. Gamma diversity in natural habitats represents the median of 1000 sampled combinations of seven and eight forest and savannah sites; error bars indicate observed range. Alpha diversity was consistently higher in farmland than in the natural habitat, whereas gamma diversity was consistently higher in natural sites than in farmland.

**Fig. 2.**
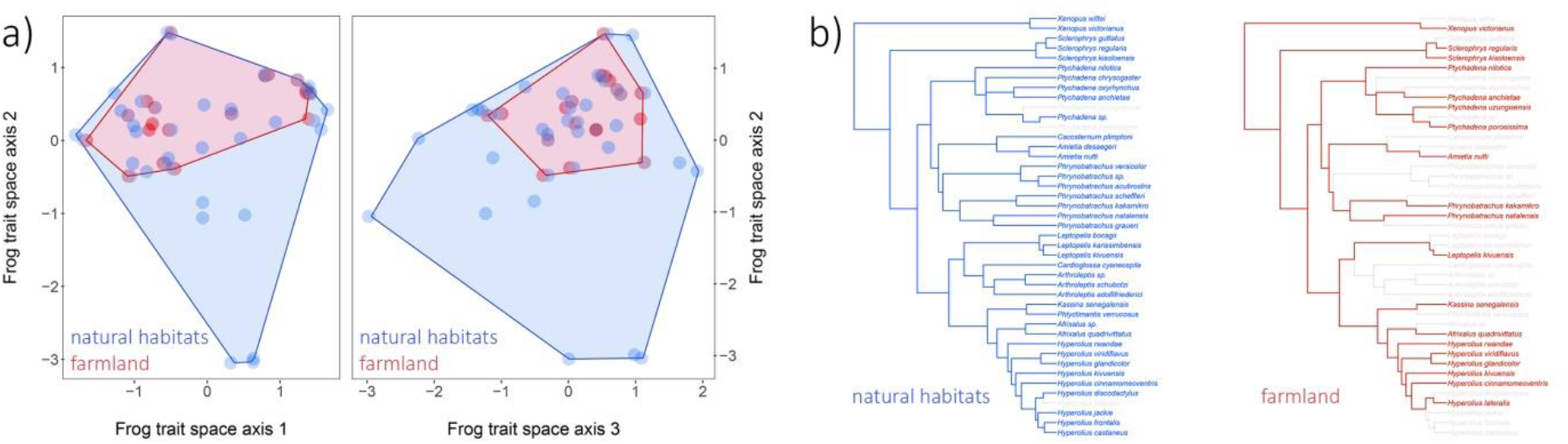
Functional and phylogenetic diversity of amphibian communities in Rwanda. a) Diversity of functional-trait combinations (functional richness) of amphibian species in natural sites (blue) vs. farmland (red) across Rwanda. Species are projected into a three-dimensional trait space where they are arranged according to the similarity in their trait combinations (left: axes 1 and 2, right: axes 2 and 3). The trait combinations of frogs found in farmland is much smaller than, and completely nested within, the trait combinations of species found in the natural sites. (b) Phylogenetic tree of the frog species found in natural sites (blue) and farmland (red) across Rwanda. While the farmland includes most of the major lineages, diversity within the lineages is greatly reduced.

The comparisons without high-elevation forest plots yielded very similar results (Appendix 2).

### Species and phylogenetic complementarity

Compared to other areas in sub-Saharan Africa, the amphibian communities in Rwanda showed moderate species richness and phylogenetic diversity (Fig. 3a,b). However, the area along the Albertine Rift (including the natural habitats in Rwanda) showed the highest species and phylogenetic complementarity, i.e. this region had the highest percentage of species and lineages with restricted ranges. In contrast, large areas in sub-Saharan Africa showed low levels of complementarity. These areas shared many widespread species and lineages and therefore showed similar taxonomic and phylogenetic composition.

**Fig. 3.**
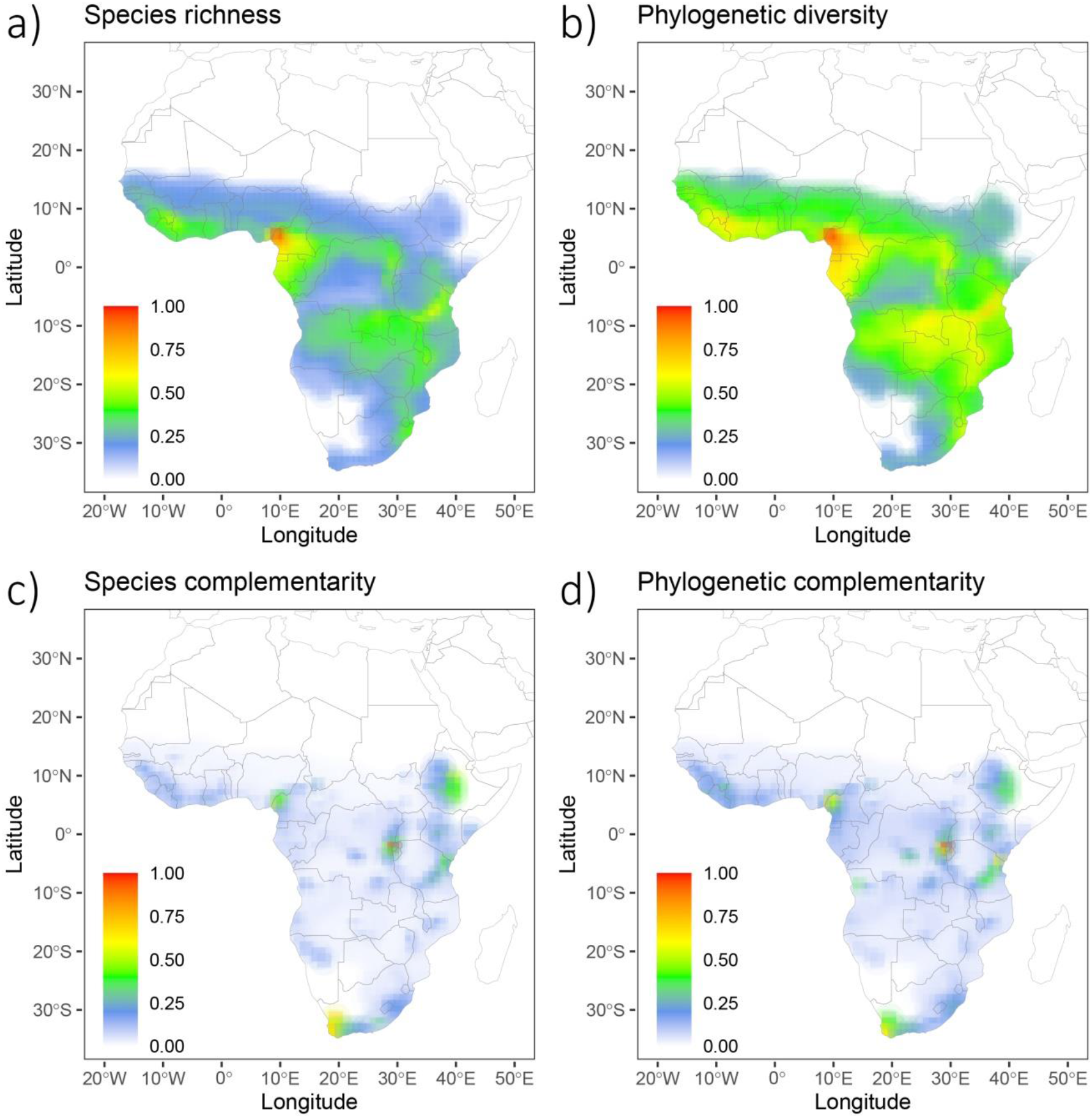
Taxonomic (species richness) and phylogenetic diversity and complementarity of amphibian species across sub-Saharan Africa at 1° resolution. All values scaled relative to the highest observed value. Species richness (a) and phylogenetic diversity (Faith’s PD; b) are calculated based the number of frog species with overlapping ranges; both peak in western Central Africa. (c) Species complementarity, the contribution of a grid cell to overall species richness relative to the number of species found in the grid cell. (d) Phylogenetic complementarity, the contribution of a grid cell to overall phylogenetic diversity relative to the diversity found in the grid cell. Grid cells that include a high percentage of species/phylogenetic lineages that are shared with few other grid cells show high complementarity; grid cells with a high percentage of species/phylogenetic lineages that occur in many other grid cells show a small complementarity. Species and phylogenetic complementarity peak in the Albertine Rift (including Rwanda), East Africa, and the Cape Region.

## Discussion

Habitat deterioration due to conversion of natural habitats into farmland drastically reduced the taxonomic, functional, and phylogenetic diversity of amphibians in Rwanda. Misleadingly, amphibian communities in farmland showed higher local alpha diversity than amphibian communities in natural habitats, which could suggest that that disturbed sites support diverse frog communities. However, communities in farmland were highly homogenised and showed almost the same species composition across the entire country. As a consequence, the regional taxonomic, functional and phylogenetic gamma diversity of amphibians in farmland across Rwanda was much lower than in natural sites, even though farmland covers a much larger area. The high alpha diversity in the farmland therefore masked the detrimental effects of habitat alterations on amphibian diversity in Rwanda (Su et al. 2004; Mori et al. 2018). This underlines that considering only local alpha diversity as an indicator for the effects of disturbance can be misleading as it ignores potential changes in species composition at both the local and the regional scale. Studies on the effect of habitat alteration on species communities should include measures of beta diversity to fully assess the impact on diversity and multifunctionality at the regional scale (Hector & Bagchi 2007; Isbell et al. 2011; Pasari et al. 2013; van der Plas et al. 2016; Mori et al. 2018, Seibold et al. 2019).

The strong decrease in functional diversity from natural forest to farmland shows that species in natural sites fulfil many unique functional roles that cannot be fulfilled by species in farmland (Gibson et al. 2011). While in some cases disturbed sites hold species communities with functional roles complementary to those in natural habitats (de Coster 2015; Riemann et al. 2017), this was not the case for amphibians in Rwanda: the functional-trait combinations of frogs from farmland were entirely nested within those from natural sites, indicating that habitat alteration only leads to a loss, not a shift, of functional roles in the amphibian communities. The homogenization of amphibian communities driven by farmland conversion hence represents a case of replacing a complex natural system with a much more simplified system (McKinney & Lockwood 1999).

The loss of taxonomic and functional diversity in farmland was partly mirrored by the loss of phylogenetic gamma diversity. Habitat alteration in Rwanda caused the loss of four highly adapted lineages with restricted geographic ranges, including the only representative of the genus *Cacosternum*, a highly adapted savannah specialist, as well as regionally endemic forest specialists with adaptations to stream breeding or direct development in the genera *Arthroleptis* and *Cardioglossa*, in the basal lineage within *Phrynobatrachus* (Zimkus et al. 2010), and in the *Hyperolius-castaneus* group (Dehling & Sinsch 2019). These species fulfill functional roles that cannot be replaced by closely-related, disturbance-tolerant species. This shows that even though the reduction of phylogenetic diversity was limited to the loss of branches within the major lineages (which implies that in disturbed sites most functions might still be maintained by closely related species), the assessment of phylogenetic diversity did not fully reflect the impact on amphibian communities revealed by the analysis of functional diversity.

The severe loss of regional taxonomic, functional, and phylogenetic gamma diversity is in line with previous studies showing that homogenization reduces regional diversity by replacing unique endemic species with widespread, abundant and often invasive species (McKinney & Lockwood 1999; Haddad & Prado 2005; Alroy 2017). The Rwandan agricultural landscape is a highly homogenous environment, which allows disturbance-tolerant amphibians to disperse without barriers (Tumushimire et al. 2020) and expand their ranges (McKinney & Lockwood 1999; McGill et al. 2015)—in fact, two of the species found in farmland (*Ptychadena porosissima*, *P*. *uzungwensis*) possibly colonized Rwanda only after the extensive modification of natural habitat (Sinsch et al. 2012; Dehling & Sinsch 2013). Amphibian species found in Rwandan farmland show common characteristics of species that thrive in human-altered environments, such as ancestral reproductive modes with high fecundity and the ability to breed throughout the year, and wide climatic niches which facilitate large geographic ranges in eastern Africa and beyond (McKinney & Lockwood 1999; Haddad & Prado 2005; Jiménez-Robles et al. 2017); none of these species is currently threatened by extinction (IUCN red list status “least concern”; IUCN 2020). In contrast, the species restricted to natural habitats show characteristics such as specialized reproduction modes, including small clutches of large eggs, direct development, highly seasonal breeding, and reproduction in streams. As a result of the widespread habitat conversion, the disturbance-tolerant species in farmland thus represent the “winners” among the Rwandan amphibians, species from natural habitats represent the “losers” (McKinney & Lockwood 1999), and the losers outnumber the winners by a wide margin. Most Rwandan amphibians from natural sites have very small geographic ranges, including several species that are endemic to the mountains of the Albertine Rift (Dehling & Sinsch 2013; Portillo et al. 2015; Channing et al. 2016; Dehling & Sinsch 2019; Fig. 3). If the remaining natural habitats of the region continue to be altered, the unique—and to a large part endemic—amphibian fauna will be lost and replaced by a small number of generalist species that are widespread across sub-Saharan Africa (Fig. 3).

Tropical ecosystems, especially in sub-Saharan Africa and South America, are expected to face even greater pressures in the future from the need to sustain a massively growing human population through expansion and intensification of agriculture (Laurance et al. 2014). Such an intensification could lead to further decline and homogenization of diversity (Karp et al. 2012; Gámez-Virués et al. 2015). However, agriculture in developing countries, including Rwanda, is still often inefficient, dominated by smallholders who lack access to modern agricultural technologies (Masters et al. 2013). The detrimental effect of habitat loss could be partly buffered if intensification of agriculture was reached via improved farming practices to close the gap between actual and potential production (Laurance et al. 2014). In addition, the introduction of alternative farming practices in semi-natural habitats, such as agroforestry systems or production forests might support more species from natural habitats than intensive agriculture (Gardner et al. 2007b; Clough et al. 2011; Edwards et al. 2014; Laurance et al. 2014), as shown for other tropical vertebrates, such as birds (Greenler & Ebersole 2015; Isbell et al. 2017; Hendershot et al. 2020). First evidence suggests that agroforestry can support high proportions of forest-associated amphibians (Angarita-M. et al. 2015; Brüning et al. 2018). Only a regional approach to biodiversity conservation will suffice to counter the detrimental effects of extensive habitat alteration and homogenization in agricultural landscapes (Seibold et al. 2019).

## Acknowledgements

Permits to enter national parks and to collect and export specimens (no. 2/ORTPN/V.U/09, 08/RDB-T&C/V.U/12, 18/RDB-T&C/V.U/12 and 10/RDB-T&C/V.U/15) were issued by the Rwandan Development Board. We thank B. Dumbo, U. Sinsch, E. Fischer, S. Seidel and H. Hinkel for help with organising and conducting field work in Rwanda. DMD acknowledges funding from the European Research Council under the European Union’s Horizon 2020 research and innovation program (grant 787638) and the Swiss National Science Foundation (grant 173342), both awarded to C. H. Graham.

## Appendix 1 Supplementary figures and tables

**Figure S1.**
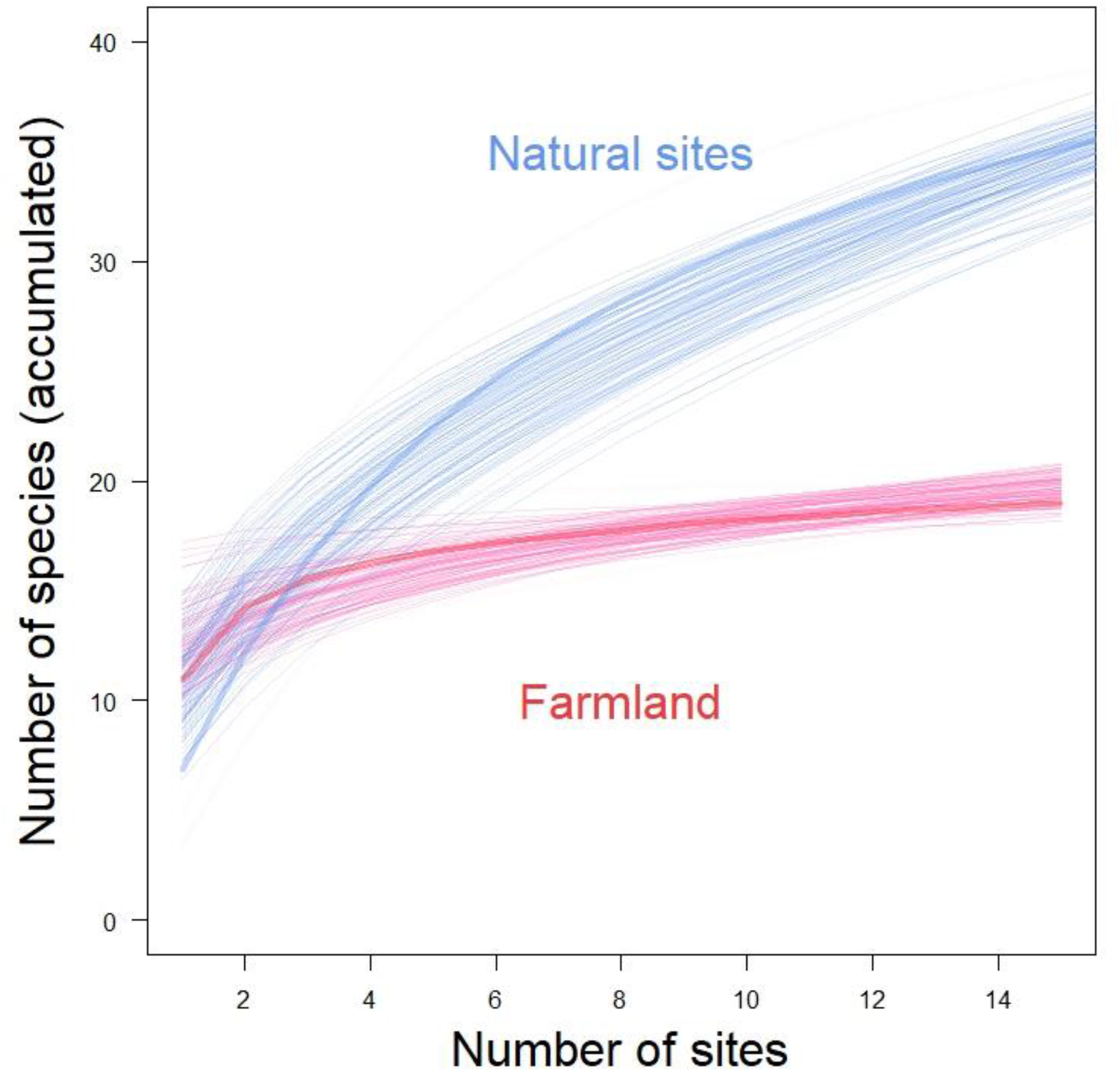
Species-accumulation curves for sites sampled in natural habitats and farmland across Rwanda. The number of species found in farmland approaches a maximum after 15 samples, whereas the number of species found in natural habitats still increases steeply.

**Table S1.**
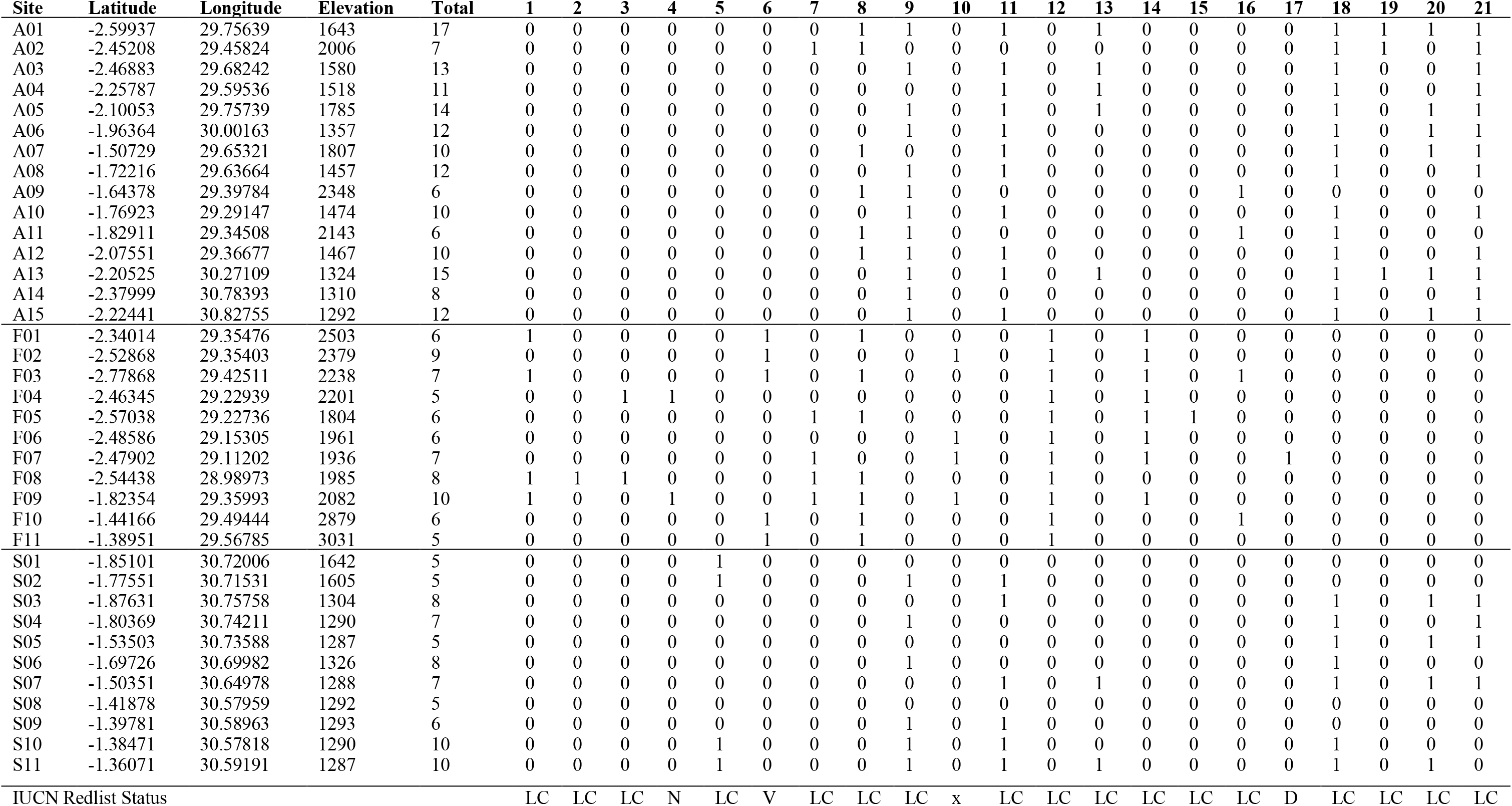
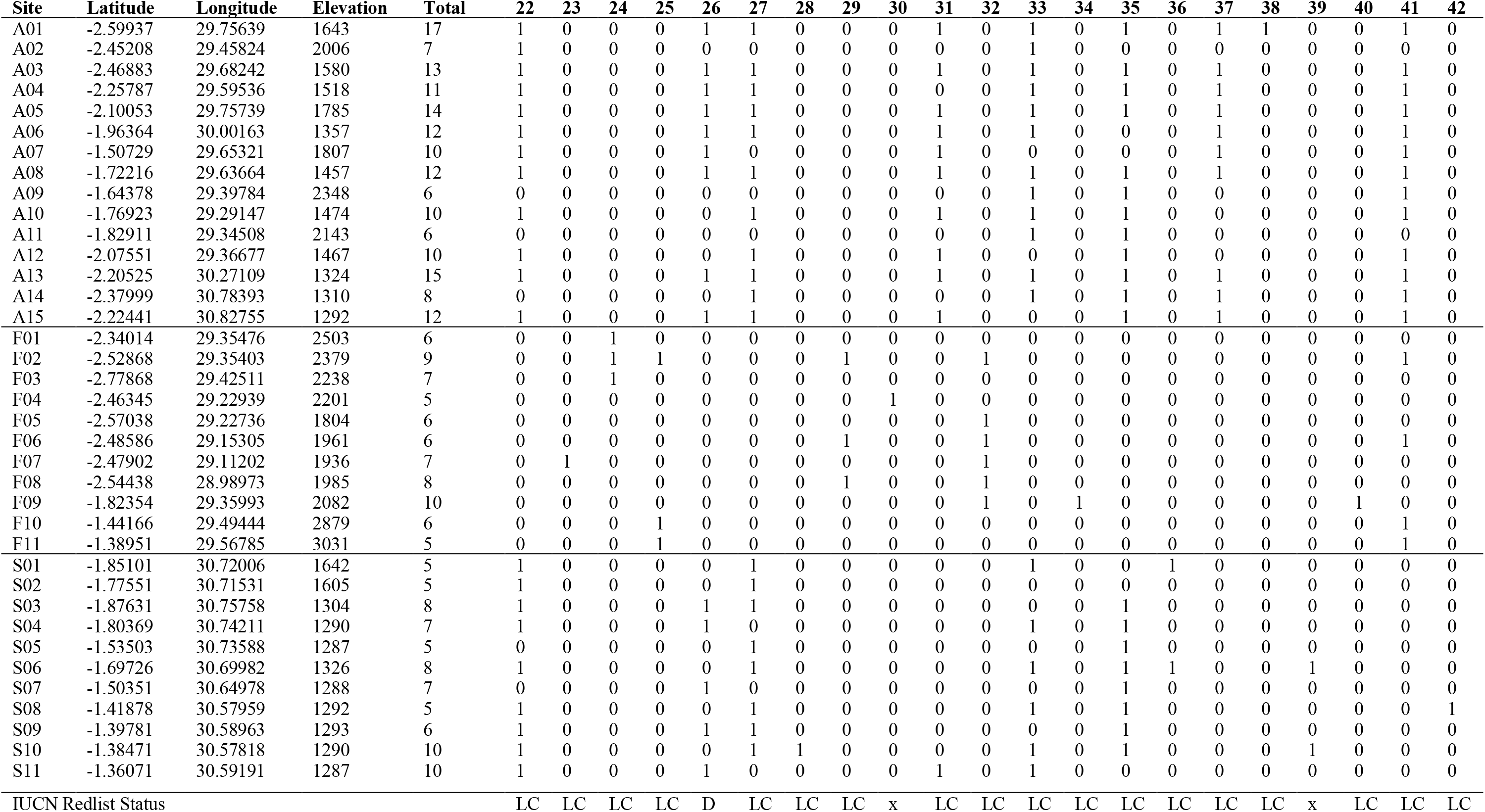
Study sites in Rwanda: latitude and longitude, elevation [m a.s.l.], total local species number and presence of species. A1–A15: agricultural marais (farmland), F01–F11: natural forest, S01–S11: natural savannah. Species identities: 1: *Arthroleptis adolfifriederici*; 2: *A. schubotzi*; 3: *Arthroleptis* sp.; 4: *Cardioglossa cyaneospila*; 5: *Leptopelis bocagii*; 6: *L. karissimbensis*; 7: *L. kivuensis*; 8: *Sclerophrys kisoloensis*; 9: *Sclerophrys gutturalis*; 10: *Afrixalus* sp.; 11: *A. quadrivittatus*; 12: *Hyperolius castaneus*; 13: *H.* cf. *cinnamomeoventris*; 14: *H. discodactylus*; 15: *H. frontalis*; 16: *H. glandicolor*; 17: *H. jackie*; 18: *H. kivuensis*; 19: *H. lateralis*; 20: *H. rwandae*; 21: *H. viridiflavus*; 22: *Kassina senegalensis*; 23: *Phlyctimantis verrucosus*; 24: *Phrynobatrachus acutirostris*; 25: *P. graueri*; 26: *P. kakamikro*; 27: *P.* aff. *natalensis*; 28: *P. scheffleri*; 29: *P. versicolor*; 30: *P. longipes*; 31: *Xenopus victorianus*; 32: *X. wittei*; 33: *Ptychadena anchietae*; 34: *Pt. chrysogaster*; 35: *Pt. nilotica*; 36: *Pt. oxyrhynchus*; 37: *Pt. porosissima*; 38: *Pt. uzungwensis*; 39: *Ptychadena* sp.; 40: *Amietia desaegeri*; 41: *A. nutti*; 42: *Cacosternum plimptoni*. Species presence: 0: absent, 1: present. IUCN (2020) redlist status of species: LC: least concern, NT: near threatened, VU: vulnerable, DD: data deficient, x: not assessed.

**Table S2.** Functional traits of frog species used in the study. The selection of traits follows Ernst et al. 2006 and Cadotte et al. 2011.

| Trait | Scale | Characteristic attributes / unit |
| --- | --- | --- |
| Basic microhabitat of postmetamorph | Nominal | 1: (semi)aquatic; 2: water edge (pond, river), on ground; 3: water edge/swamp on vegetation <2 m; 4: terrestrial incl. veg ≤ 1 m; 5: arboreal incl. tree holes; 6: semiarboreal; 7: fossorial |
| Calling site | nominal | 1: (semi)aquatic; 2: water edge (pond, river), on ground; 3: water edge/swamp on vegetation <2 m; 4: terrestrial incl. veg ≤ 1 m; 5: arboreal incl. tree holes; 6: semiarboreal; 7: fossorial |
| Egg-deposition site | nominal | 1: lentic; 2: lotic; 3: leaf litter; 4: phytotelm/rock pool; 5: burrows/underground; 6: foam nest |
| Egg-deposition type | nominal | 1: into water; 2: above water surface; 3: independent of water |
| Breeding seasonality | ordinal | 1: no marked seasonality/throughout year; 2: seasonal (at beginning of rainy season); 3: explosive (for a few days only) |
| Tadpole | nominal | 1: none; 2: free-swimming |
| Mean snout-vent length male | numeric | Mean [mm] |
| Mean head width male | numeric | Mean [mm] |
| Mean snout-vent length female | numeric | Mean [mm] |
| Mean head width female | numeric | Mean [mm] |
| Relative hind limb length | numeric | Ratio tibiofibula length/SVL |
| Hand webbing | ordinal | 1: absent or traces; 2: less than half-webbed or not between all fingers; 3: up to two-thirds webbed between all fingers; 4: nearly fully or fully webbed |
| Foot webbing | ordinal | 1: absent or traces; 2: half-webbed; 3: two-thirds-nearly fully; 4: fully webbed |
| Terminal disks | ordinal | 1: not expanded; 2: slightly expanded; 3: expanded |

Cadotte M. W., Carscadden K. & Mirotchnick N. (2011) Beyond species: functional diversity and the maintenance of ecological processes and services. – Journal of Applied Ecology 48: 1079–1087.
Ernst R., Linsenmair K. E. & Rödel M.-O. (2006) Diversity erosion beyond the species level: dramatic loss of functional diversity after selective logging in two tropical amphibian communities. – Biological Conservation 133: 143–155.

## Appendix 2

Analyses without high-elevation sites (> 2500 m).

## Results

## Comparison of alpha and gamma diversity

We recorded a total of 42 amphibian species at the study sites: 22 in forest, 17 in savannah and 19 species in farmland (Appendix 1, Table S1). Fifteen species (eleven savannah and four forest species) were shared between natural sites and farmland; three species (*Hyperolius lateralis*, *Ptychadena porosissima*, *P. uzungwensis*) were exclusively found in farmland. Alpha diversity in farmland was higher than in natural sites (*species richness*, farmland: median ± SD 11 ± 3.2, range 6–17, natural sites: 7 ± 1.9, range 4–10, Fig. S2.1a; *functional diversity*, farmland: 2.24 ± 1.01, natural sites: 1.26 ± 2.31, Fig. S2.1b; *phylogenetic diversity*, farmland: 1248 ± 292, natural sites: 799 ± 204, Fig. S2.1c). In contrast, gamma diversity was higher in natural sites than in farmland (*species richness*, natural sites 37 (range 35–39), farmland 19, Fig. S2.1a; *functional diversity*, natural sites 20.8 ± 0.73, farmland 3.9, Fig. S2.1b; *phylogenetic diversity* natural sites 2520 ± 57, farmland 1731, Fig. S2.1c).

**Figure S2.1.**
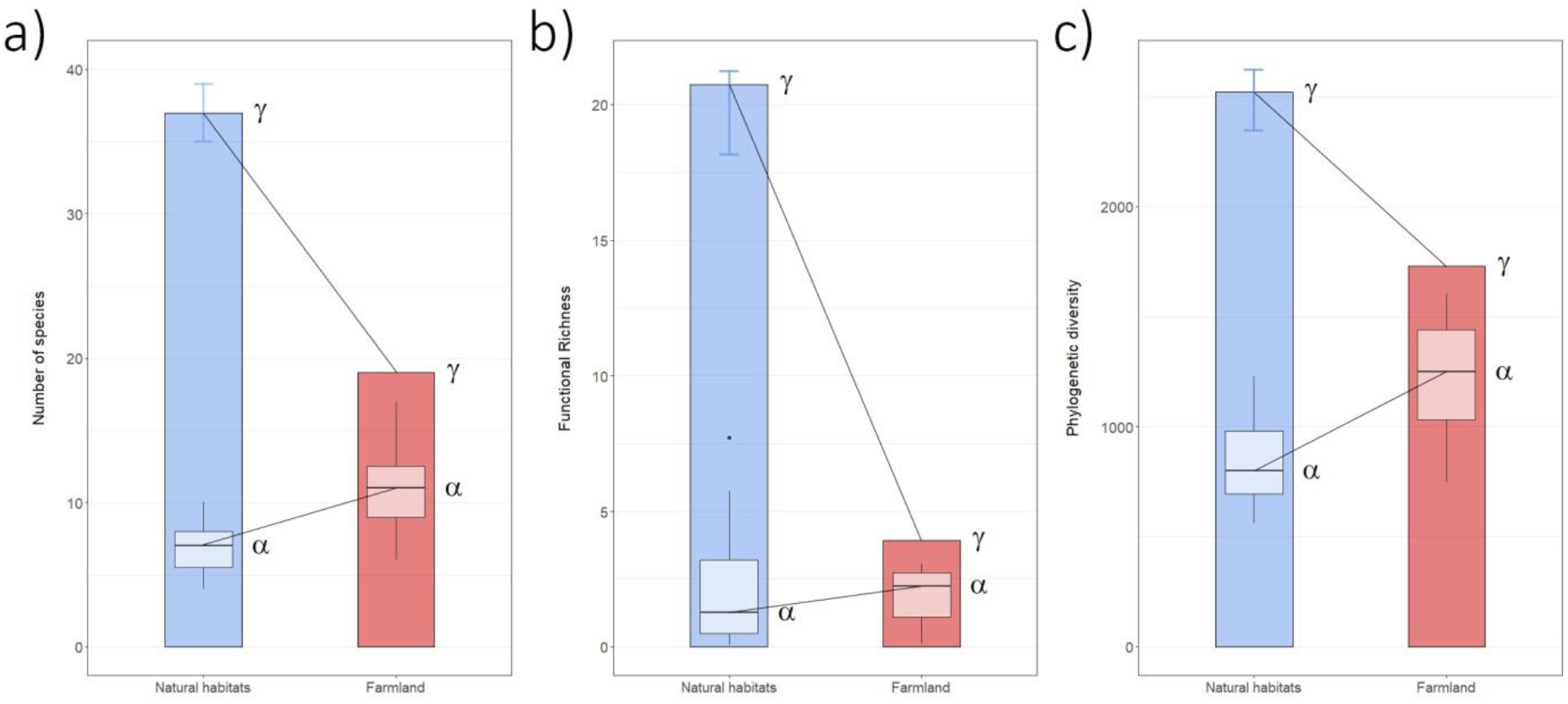
Alpha and gamma diversity of amphibian communities in natural habitats (natural forest, natural savannah; blue) vs farmland (red) across Rwanda. Diversity is measured as (a) taxonomic diversity (number of species), (b) functional diversity (functional richness), and (c) phylogenetic diversity (Faith’s PD) both on the level of local communities (alpha diversity) and once pooled across Rwanda (gamma diversity). Gamma diversity is shown as coloured bars; median and standard deviation of local alpha diversity (n = 15) are shown as lighter boxplots within bars. Gamma diversity in natural habitats represents the median of 1000 sampled combinations of seven and eight forest and savannah sites; error bars indicate observed range. Alpha diversity was consistently higher in farmland than in the natural habitat, whereas gamma diversity was consistently higher in natural sites than in farmland.

